# Coastal development and human recreation disturbance limit Australian Pied Oystercatcher *Haematopus longirostris* population sizes on E Australian beaches

**DOI:** 10.1101/2020.03.07.982223

**Authors:** Stephen Totterman

## Abstract

Australian Pied Oystercatchers *Haematopus longirostris* and their habitat were surveyed on 72 beaches and 674 km of coastline, from Fraser Island, Queensland to near the New South Wales–Victoria state border, in 2015–2018. A grand total of 232 individual birds (the sum of mean beach counts) and 41 oystercatcher territories were counted. Regression models for mean oystercatcher count density indicated a positive response to the abundance of the surf clam *Donax deltoides*, a positive New South Wales Far North Coast regional effect and a negative response to the proportion of urban beach. Models for oystercatcher territory density indicated a positive Far North Coast effect and a negative response to pedestrian access density. This report upgrades the coastal development and human recreation disturbance threats for the species.

## INTRODUCTION

The Australian Pied Oystercatcher *Haematopus longirostris* inhabits estuaries and ocean beaches along the Australian coast, with major populations in the southern states of Tasmania, South Australia and Victoria. The species is non-migratory and many breeding pairs remain on their territories throughout the year. It is also long-lived and breeding population sizes are usually limited by territorial behaviour and not by breeding success. Delayed age of first breeding is normal, resulting in a surplus of non-territorial adults in populations (see the conservation assessment by Taylor *et al*. 2014).

The Australian Pied Oystercatcher has been listed as Endangered in the state of New South Wales (NSW Government 2016) because the estimated population size is low and projected or continuing to decrease. Proposed conservation threats for the species in NSW include habitat loss, human recreation disturbance, a reduction in food resources and low breeding success, often because of depredation of eggs and chicks by the European Red Fox *Vulpes vulpes* (NSW Scientific Committee 2010).

Microtidal, high energy ocean beaches (surf beaches) are common along the SE Australian coast (Short 2007). These are essentially linear habitats with a narrow intertidal zone. Australian Pied Oystercatchers typically nest on the low front dune (Wellman *et al*. 2000). Spatial constraints and territorial behaviour mean that beach-resident oystercatcher population sizes are smaller than those counted in large estuaries (see Taylor *et al*. 2014) and are dispersed along the coast.

Commonly known as the ‘pipi’, *Donax deltoides* is a large surf clam that inhabits the intertidal and shallow subtidal zones on ocean beaches from Fraser Island, in the state of Queensland, to the Murray River, South Australia (Ferguson *et al*. 2018). Pipis are the major prey for Australian Pied Oystercatchers on ocean beaches in NSW and previous studies have reported that oystercatcher abundance is positively correlated with pipi biomass (Owner & Rohweder 2003, Harrison 2009).

Pipis are harvested by humans for food and as bait for recreational fishing. Commercial pipi fishing has been suggested to negatively impact stocks and thereby oystercatcher populations (Owner & Rohweder 2003, Harrison 2009). However, surf clams are fast-growing and short-lived, and natural variability in recruitment can also result in temporary stock declines (Murray-Jones 1999). Only hand gathering of pipis is allowed in NSW. The commercial fishery is small, with 52 licenses in 2018 and a FY 2015–16 catch of 176 t (Ferguson *et al*. 2018). There is no commercial pipi fishing in Queensland.

The feeding ecology of the Eurasian Oystercatcher *Haematopus ostralegus* and shellfishing impacts have been thoroughly studied in Europe (*e.g*. Goss-Custard 1996). However, readers are cautioned that the ecology and conservation of Australian Pied Oystercatchers on ocean beaches is unlike that for Eurasian Oystercatchers in estuaries (Owner 1997, Taylor & Taylor 2005, Harrison 2009, Taylor *et al*. 2014, Totterman 2018).

Spatial constraints amplify human recreation disturbance for shorebirds on ocean beaches. For example, mean beach widths, measured from the base of the front dune to the middle of the low tide swash zone for 11 beaches on the Far North Coast of NSW were 47–98 m (Totterman 2019a). These beach widths are similar to reported flight initiation distances for Australian Pied Oystercatchers of 39–83 m for a human stimulus (reviewed in Weston *et al*. 2012) and 129 m for a human and dog stimulus (Harrison 2009). People also disperse from beach access points and the mean along shore distance walked for 18 Far North Coast beaches was 444 m (Totterman 2019b).

This study was motivated by concerns that previous studies (Owner & Rohweder 2003, Harrison 2009) had overstated the importance of pipis for oystercatchers on ocean beaches and overlooked the coastal development and human recreation disturbance threats. The objective was to sample a larger number of beaches along the E Australian coast and re-examine oystercatcher-habitat associations.

## STUDY AREA AND METHODS

### Sampling design

The sampling space for this study extended from the Fraser Island, Queensland S to the NSW–Victoria border (Fig. 1). 117 exposed beaches with a SE–E aspect and > 1.5 km long, where pipis could be abundant (Owner 1997), were identified for the sampling frame.

**Fig. 1.**
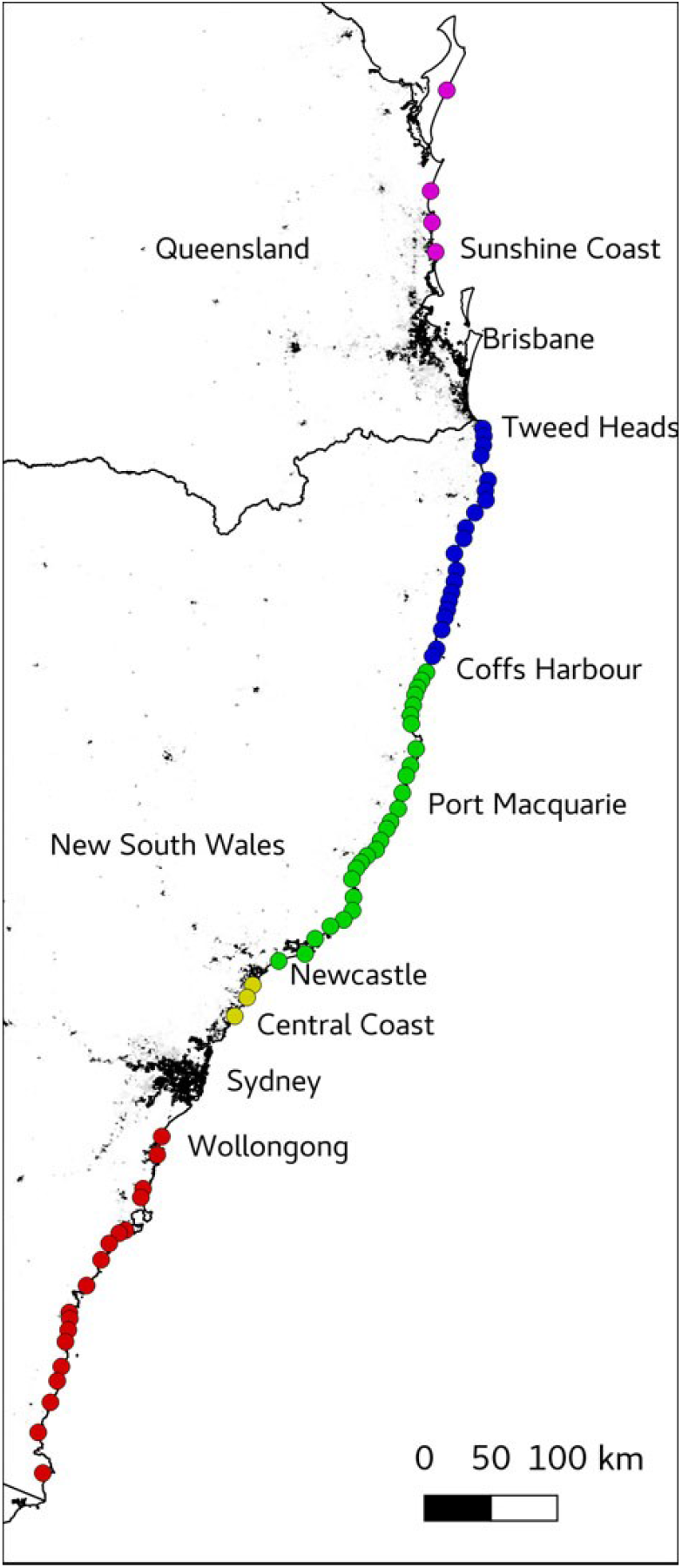
Map of the SE Australian coast showing the location of beaches (points plotted at beach centres) that were sampled in this study (Table 1). Colour-fill identifies regions: SE Queensland (magenta), Far North Coast (blue), Mid-north Coast (green), Central Coast (yellow) and South Coast (red). The background image shows human population densities (scale 0– 1000 persons/km^2^) from the Australian Population Grid 2011 (Australian Bureau of Statistics 2011). Text annotations identify states and larger coastal cities, towns and populated regions.

Beaches were defined by natural breaks, *e*.*g*. headlands and rivers. Some short beaches were combined where the break was a minor obstacle, *e*.*g*. a short, low rocky interval that could easily be traversed at all tides. Sampling prioritised longer, less-developed beaches, where oystercatchers could be abundant (Owner 1997, Harrison 2009), for the purpose of estimating bird-habitat models (Albert *et al*. 2010). Selection of high pipi and/or oystercatcher abundance beaches was also guided by other studies and reports (Gray 2016a,b, Totterman 2019a, NSW Department of Planning, Industry and Environment (DPIE) 2019a).

**Table 1.**
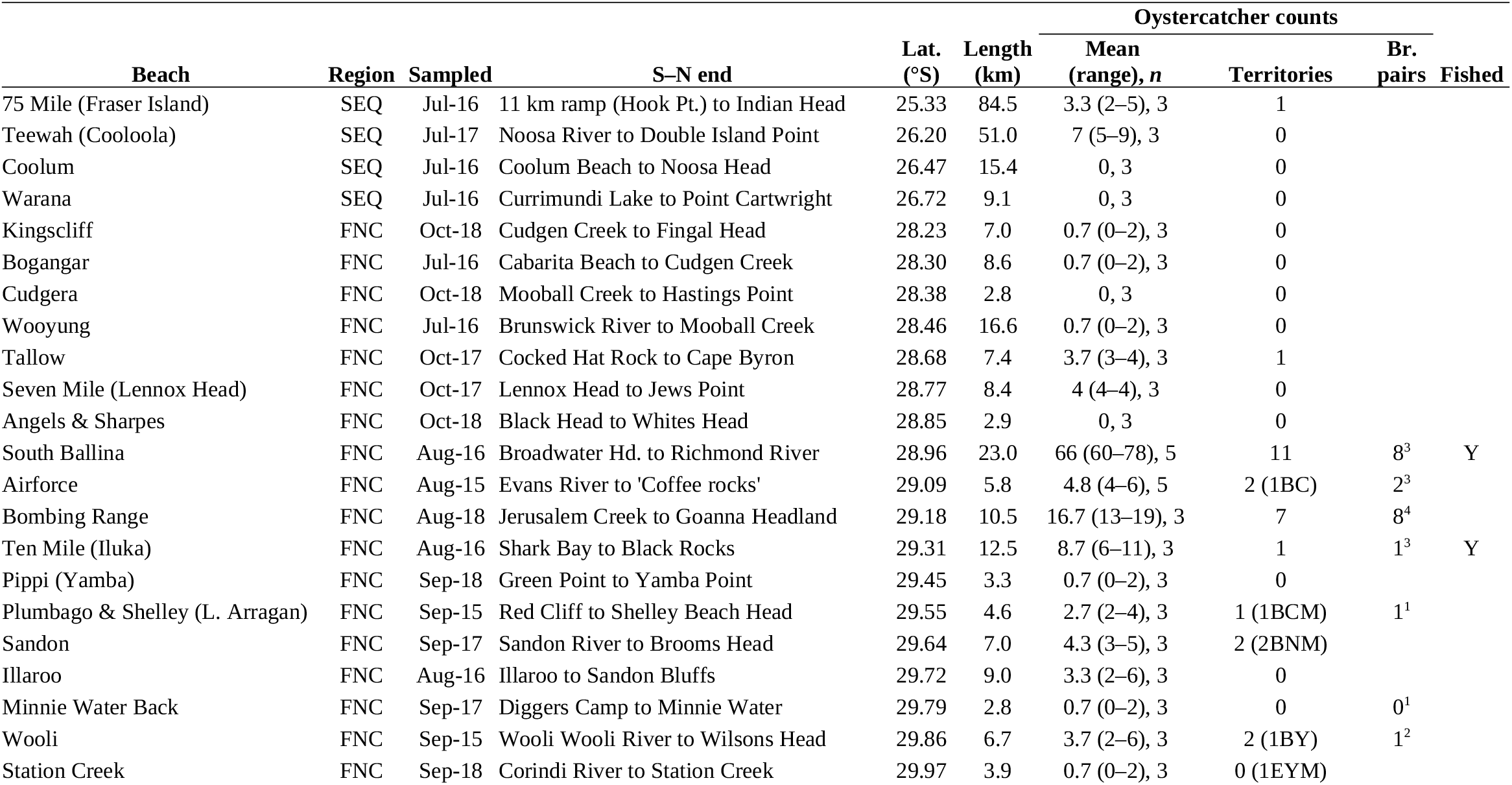
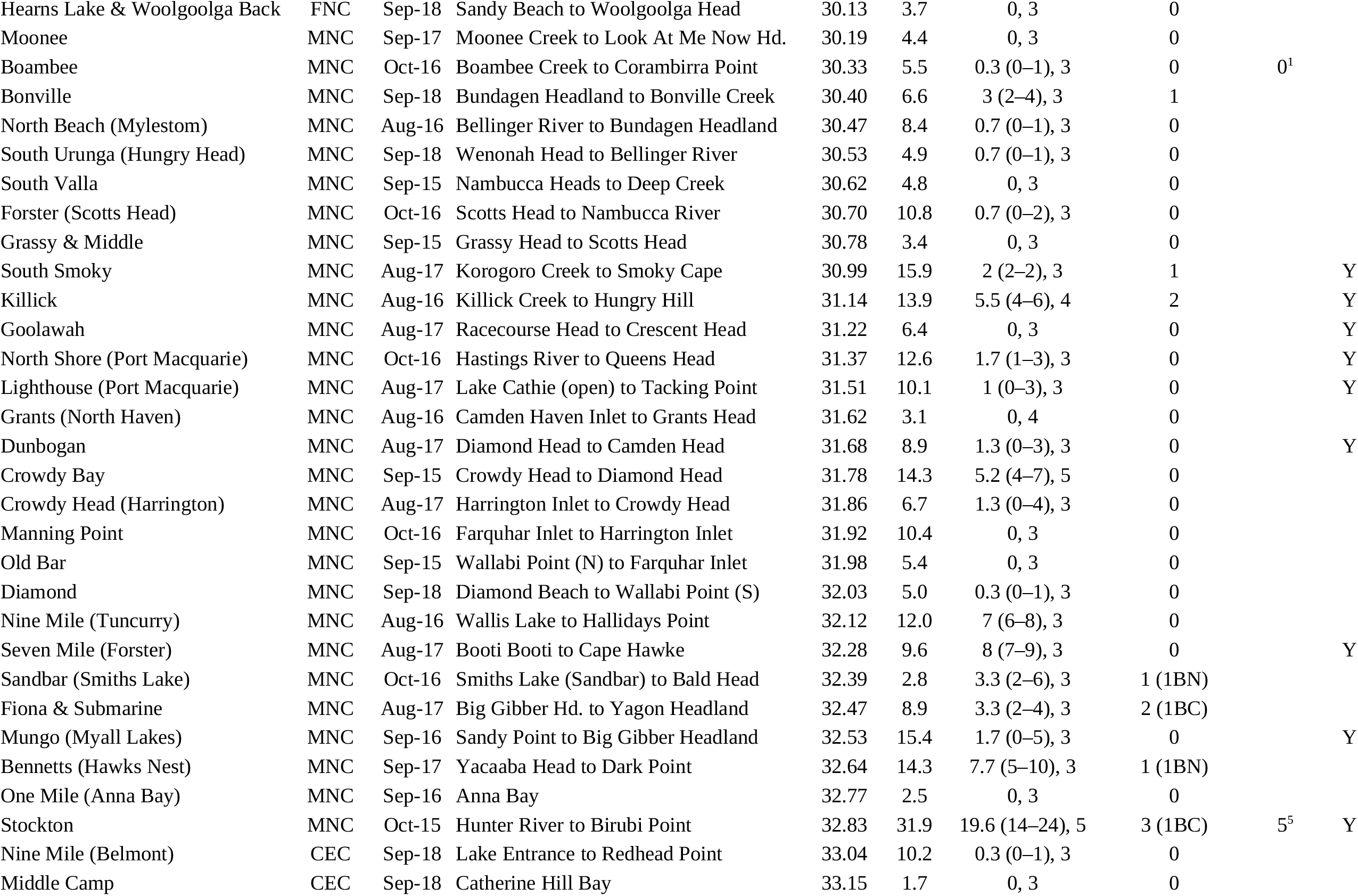
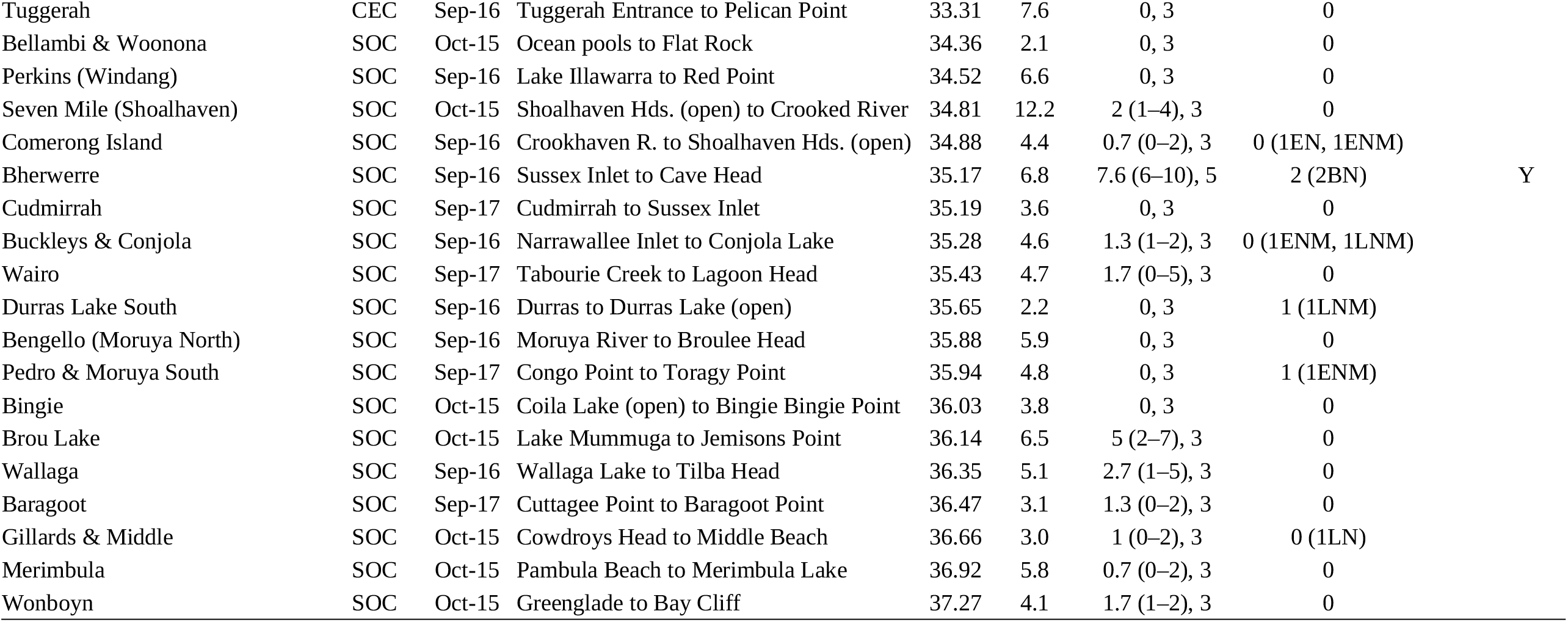
Beaches that were sampled in this study and oystercatcher counts (beaches are ordered N to S by mid-point latitude). Regions are SE Queensland (SEQ), the Far North Coast of NSW (FNC), Mid-north Coast (MNC), Central Coast (CEC) and South Coast (SOC). Coded breeding records accompany beach territory counts: *n* pairs; habitat (Beach, Estuary, Lake); observation (C = Copulation; N = nest with eggs; Y = dependent young), and; M indicates marked nests (*e*.*g*. with fencing and signage). For example, 2BNM translates to 2 Beach Nests Marked. Several of these observations were of estuary/lake residents and not beach territories. Breeding pair count data sources were: 1) Harrison 2009 (br. season 2005); 2) Anon., pers. comm. 2015; 3) NSW DPIE unpubl. data (br. season 2016), and; 4) NSW DPIE unpubl. data (br. season 2018), and; 5) Fraser & Lindsey 2018 (br. seasons 2014–2017; there were only five reliable territories in that study and surely < five breeding pairs). Nest coordinates were used to assign breeding pairs from the NSW DPIE reports to beaches defined in this study. Beaches with commercial pipi fishing are indicated (Y).

The oystercatcher breeding season on the Far North Coast of NSW is Aug–Dec and late attempts generally are replacement clutches (Wellman *et al*. 2000). Beaches were sampled in Jul–Oct, early in the breeding season, when competition for territories and territory occupancy was assumed to be maximum. Sampling started with beaches in the N, progressed ‘down’ the coast and included a few beaches on the return trip. Sampling in any one year selected non-adjacent beaches to reduce spatial autocorrelation in oystercatcher counts that could result from short-distance movements.

### Counts

Beaches were surveyed with a handheld Global Positioning System satellite receiver (Garmin GPS 76), marking 4–13 (mode 7) regularly spaced along shore waypoints (*i*.*e*. coordinates saved in the device) that were used to measure beach length and distances for pipi sampling and oystercatcher territory mapping.

Oystercatchers were usually counted at low tide and early in the morning, when there were fewer people on beaches that could disturb birds and displace them along shore. A four-wheel-drive (4WD) vehicle was used to count longer beaches, where permitted. Otherwise, and for shorter beaches, a bicycle was used. Some of the shortest beaches, and those with a steep face and soft sand, were counted on foot.

Oystercatchers present in any estuaries, lakes and associated sand spits adjacent to a beach were noted but not added to the beach count. For example, coastal lakes are common along the NSW South Coast, many of these were inhabited by oystercatchers and those birds can be found roosting and sometimes nesting near the beach. However, those birds forage and spend most of the time in the lakes.

‘Adult’-plumage oystercatchers were counted including any second year immature plumage birds which superficially look the same as adults (Marchant & Higgins 1993). More precise age classification was not always possible (with variable observation distances, light, for birds in flight, etc.) and beaches had to be counted quickly to avoid along shore movements of birds. Any birds that were marked with bands or alphanumeric leg flags were noted to assist in the identification of local residents.

Territory mapping (spot mapping) can easily be applied to beach-resident oystercatchers because they are large, conspicuous birds and territories on ocean beaches are practically one-dimensional (Totterman 2018). The minimum territory mapping effort was three counts per beach and on non-consecutive days, to reduce serial correlation. Beaches with higher oystercatcher counts also tended to have higher territory counts. Effort was increased to ≥ five counts for beaches with mean oystercatcher count densities > 0.5 birds/km, based on breeding season densities of 0.3–1.9 birds/km for beaches on the Far North Coast of NSW (Owner & Rohweder 2003, Harrison 2009). Additional, opportunistic counts were also made for some beaches where pipi sampling continued beyond the initial counting period. Territories were identified as adult ‘pairs’ that were present on the same stretch of beach on ≥ 50% of counts (*e*.*g*. 2/3 or 3/5). These are rapid, low-cost estimates and not all pairs that occupy a stretch of beach may breed. Territories are compared with independent counts of breeding pairs in the Results.

### Pipi abundance

A new ‘feet digging’ method (Totterman 2019c) was used to efficiently sample pipis and enable precise beach-scale estimates of pipi abundance and mean pipi length. The method was developed from a recreational fishing technique that involves twisting one’s feet into the thixotropic sand to dislodge buried clams which are then recovered by hand. Several plots are sampled across the swash zone in one 5-min sampling unit and the process is replicated at several locations along the beach. Mean feet digging pipi counts have been shown to correlate very strongly with quadrat-based mean transect pipi densities (*r* = 0.98) and pipi length-frequency distributions were similar to those from quadrat-based samples except that feet digging is not effective for pipis < 16 mm (Totterman 2019c). These young pipis are not always present or abundant and can be ignored because individual mass increases exponentially with shell length (Murray-Jones 1999) and large clams contribute more to oystercatcher diets (Totterman 2018).

Variance increases strongly with mean pipi counts (Totterman 2019c). A constant precision stopping rule was used to prevent oversampling of low pipi abundance beaches and undersampling of high abundance beaches. A first pass sampled pipis at 20 locations, regularly spaced along shore from a random starting distance and cycling through the tidal stages high, ebb, low and flow (*i*.*e*. four tidal stages × five locations per tidal stage × 5 minutes per location). If the SE of the mean pipi count was > 2 pipis/5 min, then further sampling passes, again with 20 locations in each pass, were performed until the SE was *≤* 2 pipis/5 mins or the cumulative sample size was 100. Stratified sampling was applied to three long beaches, Teewah, Bennetts and Stockton (Table 1), which each had a distinct low pipi abundance section at the southern end.

Pipi lengths were measured with a caliper to the nearest mm (maximum shell length). A maximum sample size stopping rule was used to save time on high pipi abundance beaches. All pipis were measured in the first sampling pass (see above). If pipi abundance was low and the mean count from the first pass was precise (SE *≤* 2 pipis/5 mins), then mean length was based on < 100 pipis. Otherwise, and if < 100 pipis were sampled in the first pass, pipis were also measured in each subsequent pass until the sample size was ≥ 100, which was large enough for a precise estimate of mean pipi length. After that, pipis were only counted.

### Human recreation and other habitat variables

People, dogs, 4WDs, motorcycles, bicycles, horses and, on Fraser Island, aircraft were counted simultaneous with oystercatchers. Dogs included dingos (naturalised dogs) that were present in low densities on Fraser Island and in the Myall Lakes region of NSW. Counts of people and dogs excluded those in 4WDs or in the surf. Motorcycle, bicycle, horse and aircraft counts were small, with grand totals of three, 32, 42 and four respectively, and were not included in the statistical analysis.

Coastal development variables and indirect measures of human recreation measured were: urban length, length of public beach driving zones, whether or not beach driving permits were required, length of dog exercise zones, length of beach adjoining any terrestrial protected areas (National Parks and Nature Reserves), counts of pedestrian and, where public beach-driving was permitted, 4WD beach access tracks and counts of beach front and near-beach camping grounds and caravan parks. Access tracks include public and private, formal and informal tracks. Tracks had to be frequently used and lead somewhere to be counted, *e*.*g*. temporary dune driving tracks were not counted. 4WD tracks were also used by pedestrians on many beaches but were not added to that count. There was no public beach driving for the majority of beaches sampled and so the driving permits variable was not included in the statistical analysis.

Human population density was calculated from the 1 × 1 km Australian Population Grid 2011 (Australian Bureau of Statistics 2011). The intersection of a line defined by GPS survey waypoints with the Australian Population Grid 2011 identifies beach front grid cells and the sum of these cells equals the ‘beach front’ human population.

Beach morphodynamic state (Short 2007) was observed during moderate surf conditions and one handful of intertidal sand was sampled at each GPS waypoint. Sand grain size was classified on the Wentworth scale by visual comparison to graded samples on a sand gauge. Any shell fragments present were ignored. Beach morphodynamic state and sand grain size within beaches were summarised as modal values. Sixty-two beaches sampled (86%) had transverse bar and rip inner bars and 64 (89%) had medium sand (0.25–0.5 mm). These are the most common conditions for ocean beaches in NSW (Short 2007). With such little variation, these two variables were not included in the statistical analysis.

### Statistical analysis

There were two response variables: mean oystercatcher count and territory count, and 19 input variables (Table 2). Input variables are all of those measured, while predictors are the terms that are actually used in statistical modelling (Gelman & Hill 2007). Variables that scale with beach length were converted to linear densities (*e*.*g*. birds/km) and proportions (*e*.*g*. proportion of urban beach).

**Table 2.** Variables used in the statistical analysis. ‘Extensive variables’ were converted to densities (*dens*.) and proportions (*prop*.) by dividing by beach length. Predictors identified in the fifth column (B for mean oystercatcher count models and T for oystercatcher territory count models) are those remaining after data exploration and reduction (see Results).

| Description | Measured | Scale | Predictor | Model |
| --- | --- | --- | --- | --- |
| **Response variables** |  |  |  |  |
| Oyc. mean count | *count.B* | length | *dens.B* |  |
| Oyc. terr. count | *count.T* | length | *dens.T* |  |
| **Input variables** |  |  |  |  |
| Beach length (km) | *length* |  | *length* | B, T |
| Urban length (km) | *length.urban* | length | *prop.urban* | B, T |
| 4WD length (km) | *length.4WD* | length | *prop.4WD* | B, T |
| Dogs length (km) | *length.dogs* | length | *prop.dogs* | B |
| Dogs off-leash length (km) | *length.offleash* | length | *prop.offleash* | B, T |
| Protected area length (km) | *length.protect* | length | *prop.protect* | B |
| Pedestrian access count | *access.people* | length | *dens.acc.people* | B, T |
| 4WD access count | *access.4WD* | length | *dens.acc.4WD* |  |
| Camping & c’van parks count | *count.camps* | length | *dens.camps* | B |
| People mean count | *count.people* | length | *dens.people* |  |
| Dogs mean count | *count.dogs* | length | *dens.dogs* |  |
| 4WDs mean count | *count.4WDs* | length | *dens.4WDs* |  |
| Human population (/km<sup>2</sup>) | *pop1* | length×100 | *dens.pop1* (/Ha) | B, T |
| Mean pipis count (/5 min.) | *pipis* |  | *pipis* | B, T |
| Pipi mean length (mm) | *pipis.length* |  | *pipis.length* | B, T |
| UTM x-coordinate (m) | *x* | 1000 | *x* (km) |  |
| UTM y-coordinate (m) | *y* | 1000 | *y* (km) |  |
| Commercial pipi fishing | *fished* (Y/N) |  | *fished* | B, T |
| Region | *region*<br>(SEQ/FNC/<br>MNC/CEC/SOC) |  | *region2*<br>(FNC/Other) | B, T |

Exploratory data analysis was performed to detect patterns, multicollinearity and to reduce the number of predictors. This involved three steps: 1) spatial patterns (‘bubble maps’, correlograms); 2) bivariate associations and interactions (scatter plots), 3) variable clustering, and; 4) variance inflation factors (to identify multicollinearity).

A spline correlogram (Bjørnstad & Falck 2001) for mean oystercatcher count density indicated a spatial autocorrelation range of 63 km (95% CI 28–111 km). A radius of 50 km was used for calculating binary spatial weights for spatial statistics.

Multiple regression models were fitted to oystercatcher counts because they allow for simple and interpretable inference and the sample size was unbalanced and effectively small, with high oystercatcher counts on only a few beaches. Repeated-measures models were not used because mean oystercatcher counts were fairly precise, the number of counts per beach was as small as three and estimating the variance among repeated counts was of no interest (Murtaugh 2007).

Multicollinearity seriously confounds the interpretation of multiple regression results (Graham 2003, Cade 2015). The first step to reducing multicollinearity was a cluster analysis of bivariate correlations among continuous input variables. The dendrogram was cut at Spearman *r*^2^ = 0.49 (|*r*| = 0.7) and one representative variable was selected for each cluster (Dormann *et al*. 2013). Next, variance inflation factors (VIFs) were computed for multiple regression models with the reduced set of variables from the cluster analysis. Variables with large VIFs were dropped backward-stepwise until all VIFs were < 3 (Zuur *et al*. 2010). To avoid dropping useful predictors, variable selection in both of these analyses was made with consideration to the strength of bivariate associations with oystercatcher count densities and practical and *a priori* reasons (Dormann *et al*. 2013).

Generalised linear models (GLMs) were fitted to oystercatcher counts with a log-link and log(beach length) as an offset, to effectively model count density. A negative binomial error distribution was assumed for integerised mean oystercatcher counts (rounded to zero decimal places) and a Poisson distribution was assumed for territory counts. GLM assumptions were checked using overdispersion statistics, expected zeros (Warton 2005) mean-residual variance plots (Ver Hoef & Boveng 2007) and diagnostic plots using Dunn-Smyth randomised quantile residuals (Dunn & Smyth 1996).

Model selection was automated with three steps: 1) an exhaustive search of all subsets of models, ranking them by small sample Akaike’s Information Criterion (AICc); 2) selecting the ΔAICc < 6 model set, and; 3) removal of models from the ΔAICc < 6 set that where more complex versions of those having a lower AICc value (the ‘nesting rule’ of Richards 2008, also see Arnold 2010).

Null models fitted to oystercatcher counts included the predictor mean pipi count with a region covariate (Owner & Rohweder 2003). The pipis:region interaction was also included in the exhaustive search of all subsets of models and marginality constraints were respected, *i*.*e*. main effects were always included together with the interaction term. ‘Nil models’ for oystercatcher counts were intercept-only models, *i*.*e*. with no predictor variables. The nil model intercept is the logarithm of the grand mean oystercatcher count density.

Goodness of fit for oystercatcher count models was examined by plotting fitted versus observed densities and calculating likelihood ratio pseudo-*R*^2^ (Nagelkerke 1991) with reference to nil models. Predictive performance (*e*.*g*. cross-validation accuracy) was not evaluated because there were few high density sites for testing.

All GIS and statistical analyses were performed using the software R version 4.0.4 (R Core Team 2021). Geographic analyses used the R packages *sp* version 1.4.5 (Bivand *et al*. 2013), *rgdal* version 1.5-23 (Bivand *et al*. 2021) and *raster* version 3.4-5 (Hijmans 2020). Spatial statistics used *ncf* version 1.2-9 (Bjørnstad 2020) and *spdep* version 1.1-5 (Bivand *et al*. 2013). Clustering of bivariate correlation coefficients was performed using *Hmisc* version 4.5-0 (Harrell 2021). Variance inflation factors were computed using *car* version 3.0-10 (Fox & Weisberg 2019). Negative binomial models were fitted using *MASS* version 7.3-53 (Venables & Ripley 2002). Randomized quantile residuals were computed using *statmod* version 1.4.35 (Dunn & Smyth 1996). Model searching, ranking and selection was performed using the functions dredge and nested in *MuMIn* version 1.43.17 (Barton 2020). Spatial autocorrelation in GLM residuals was checked using the lm.morantest function in *spdep*. While this function is not intended for GLMs, it could be useful for relative comparisons among models and there were no ready alternatives that take into account predictor variables.

## RESULTS

### Spatial patterns

72 of 117 beaches (62%) from the sampling frame and 674 of 942 km (72%) were sampled including all of the longer beaches (except for Budgewoi on the Central Coast of NSW), all of the known high pipi abundance beaches and all beach Key Management Sites for the Australian Pied Oystercatcher (Fig. 1). The coastline was divided into five *ad hoc* contiguous regions: SE Queensland (from Fraser Island S to the Queensland-NSW border); Far North Coast (to Coffs Harbour); Mid-north Coast (to the Hunter River); Central Coast (to Broken Bay), Sydney (to Port Hacking), and; South Coast (to the NSW-Victoria border). Sample sizes were larger for regions where oystercatchers were more abundant: 20 of 28 beaches from the Far North Coast; 27 of 30 from the Mid-north Coast, and; 18 of 37 from the South Coast. Three of seven beaches from the sampling frame were sampled from the Central Coast (where only one oystercatcher was counted), zero of four beaches from Sydney and four of 11 beaches from SE Queensland.

A grand total of 232 individual Australian Pied Oystercatchers (the sum of mean counts) and 41 territories were counted. Narrow count ranges (Table 1) indicate that there were no substantial movements of birds in to and out of beaches between counts. Territory estimates agreed reasonably well with independent breeding pair counts (Fig. 2).

**Fig. 2.**
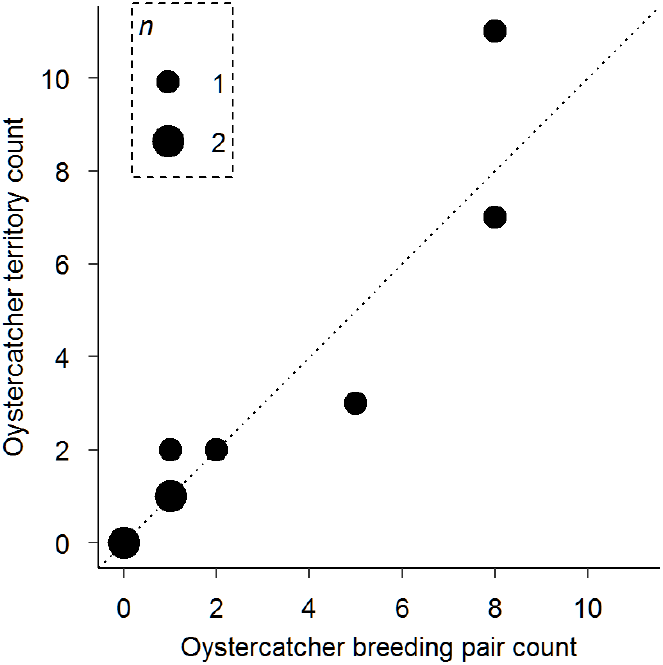
Oystercatcher territory estimates compared to breeding pairs (Table 1). Point sizes are proportional to the number of observations. A statistical comparison is not useful because the sample size (nine beaches) was small and there is insufficient power to detect small deviations from the 1:1 line of agreement (dotted line).

The spatial pattern for mean oystercatcher count densities was strongly aggregated with a ‘hot spot’ in the Richmond River area of the Far North Coast (Figs. 3a,b). Mean pipi counts showed a different spatial pattern, with a hot spot in the Mid-north Coast region (Fig. 3c). Large pipis were available on all beaches. Mean pipi length increased from 37 mm on the South Coast to 46 mm on the Far North Coast (Fig. 3d).

**Fig. 3.**
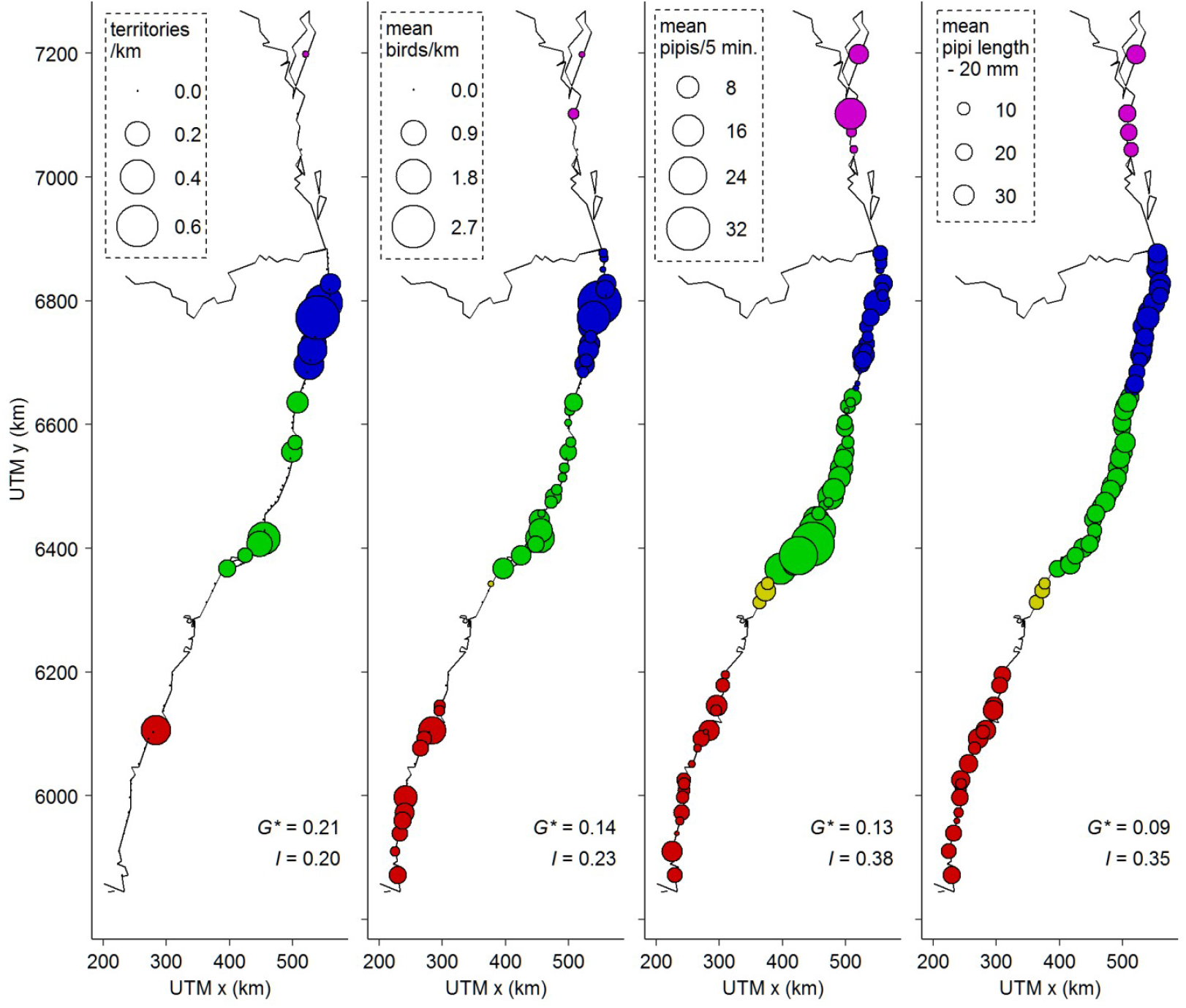
Oystercatcher and pipi spatial patterns along the E Australian coast. Point sizes are proportional to abundance/size and colour identifies regions: SE Queensland (magenta), Far North Coast (blue), Mid-north Coast (green), Central Coast (yellow) and South Coast (red). Note that mean pipi length is offset by 20 mm. Getis-Ord *G\** and Moran’s *I* statistics measure spatial concentration and similarity (autocorrelation) respectively for which all *P*-values were < 0.001.

### Bivariate associations

Mean oystercatcher count density was positively correlated with mean pipi count and the proportion of protected beach (Spearman *r* ≥ 0.38) and negatively correlated with the proportion of urban beach, human population density, mean people count density, mean dog count density, pedestrian access count density and the proportion of dogs permitted beach (Spearman *r* ≤ −0.39) (Table 3, Fig. 4).

**Table 3.** Spearman rank-based correlation matrix for all continuous input variables (Table 2). Weak correlations (|*r*| < 0.3) are not presented.

|  | dens.B | length | prop.urban | prop.4WD | prop.dogs | prop.offleash | prop.protect | dens.access.people | dens.access.4WD | dens.camps | dens.people | dens.dogs | dens.4WD | dens.pop1 | pipis | pipis.length | x | y |
| --- | --- | --- | --- | --- | --- | --- | --- | --- | --- | --- | --- | --- | --- | --- | --- | --- | --- | --- |
| dens.T | 0.6 |  |  |  |  |  | 0.4 | -0.4 |  |  |  |  |  |  |  | 0.3 |  |  |
| dens.B |  |  | -0.5 |  | -0.4 |  | 0.5 | -0.5 |  |  | -0.4 | -0.5 |  | -0.5 | 0.4 |  |  |  |
| length |  |  |  | 0.5 |  |  |  | -0.4 | 0.3 |  |  |  | 0.3 |  | 0.5 | 0.3 |  | 0.3 |
| prop.urban |  |  |  |  | 0.4 | 0.4 | -0.6 | 0.6 |  |  | 0.8 | 0.8 |  | 0.8 |  |  |  |  |
| prop.4WD |  |  |  |  |  |  |  | -0.4 | 0.9 |  |  |  | 0.8 |  | 0.3 | 0.3 |  | 0.3 |
| prop.dogs |  |  |  |  |  | 0.5 | -0.7 | 0.4 |  |  | 0.4 | 0.5 |  | 0.5 |  |  |  |  |
| prop.offleash |  |  |  |  |  |  | -0.6 |  |  |  |  | 0.4 |  | 0.3 |  |  |  |  |
| prop.protect |  |  |  |  |  |  |  | -0.6 |  |  | -0.6 | -0.7 |  | -0.5 |  |  |  |  |
| dens.access.people |  |  |  |  |  |  |  |  | -0.3 |  | 0.7 | 0.5 |  | 0.6 |  |  |  |  |
| dens.access.4WD |  |  |  |  |  |  |  |  |  |  |  |  | 0.8 |  | 0.3 |  |  |  |
| dens.camps |  |  |  |  |  |  |  |  |  |  |  |  |  |  |  |  |  |  |
| dens.people |  |  |  |  |  |  |  |  |  |  |  | 0.8 |  | 0.7 |  |  |  |  |
| dens.dogs |  |  |  |  |  |  |  |  |  |  |  |  |  | 0.7 |  |  |  |  |
| dens.4WD |  |  |  |  |  |  |  |  |  |  |  |  |  |  | 0.3 |  |  |  |
| dens.pop1 |  |  |  |  |  |  |  |  |  |  |  |  |  |  |  |  |  |  |
| pipis |  |  |  |  |  |  |  |  |  |  |  |  |  |  |  |  |  |  |
| pipis.length |  |  |  |  |  |  |  |  |  |  |  |  |  |  |  |  | 0.5 | 0.5 |
| x |  |  |  |  |  |  |  |  |  |  |  |  |  |  |  |  |  | 1.0 |

**Fig. 4.**
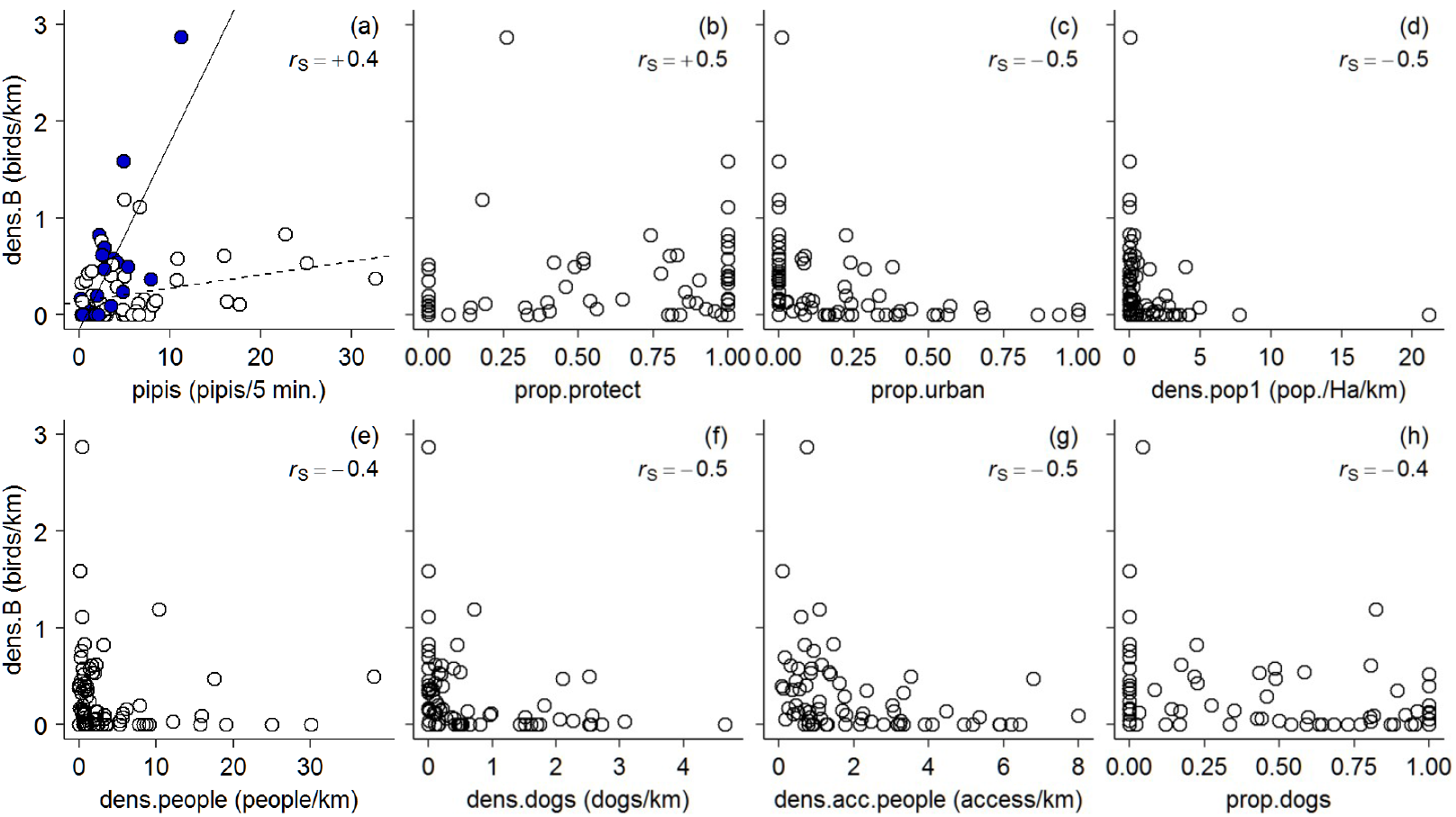
Mean oystercatcher count density correlations. Variables are described in Table 2 and Spearman rank-based correlations are in Table 3. Far North Coast beaches (Coffs Harbour to the NSW-Queensland border) are plotted as blue-filled circles and other regions are plotted as white-filled circles in (a). Oystercatcher-pipi lines in were fitted by ordinary least squares linear regression with *region2* (Far North Coast/Other) as a covariate, however the reported correlation statistic does not include that covariate. These notes also apply to Fig. 5.

Oystercatcher territory count density was positively correlated with mean pipi count, mean pipi length and the proportion of protected beach (Spearman *r* ≥ 0.28) and negatively correlated with pedestrian access count density and human population density (Spearman *r* ≤ −0.30) (Table 3, Fig. 5).

**Fig. 5.**
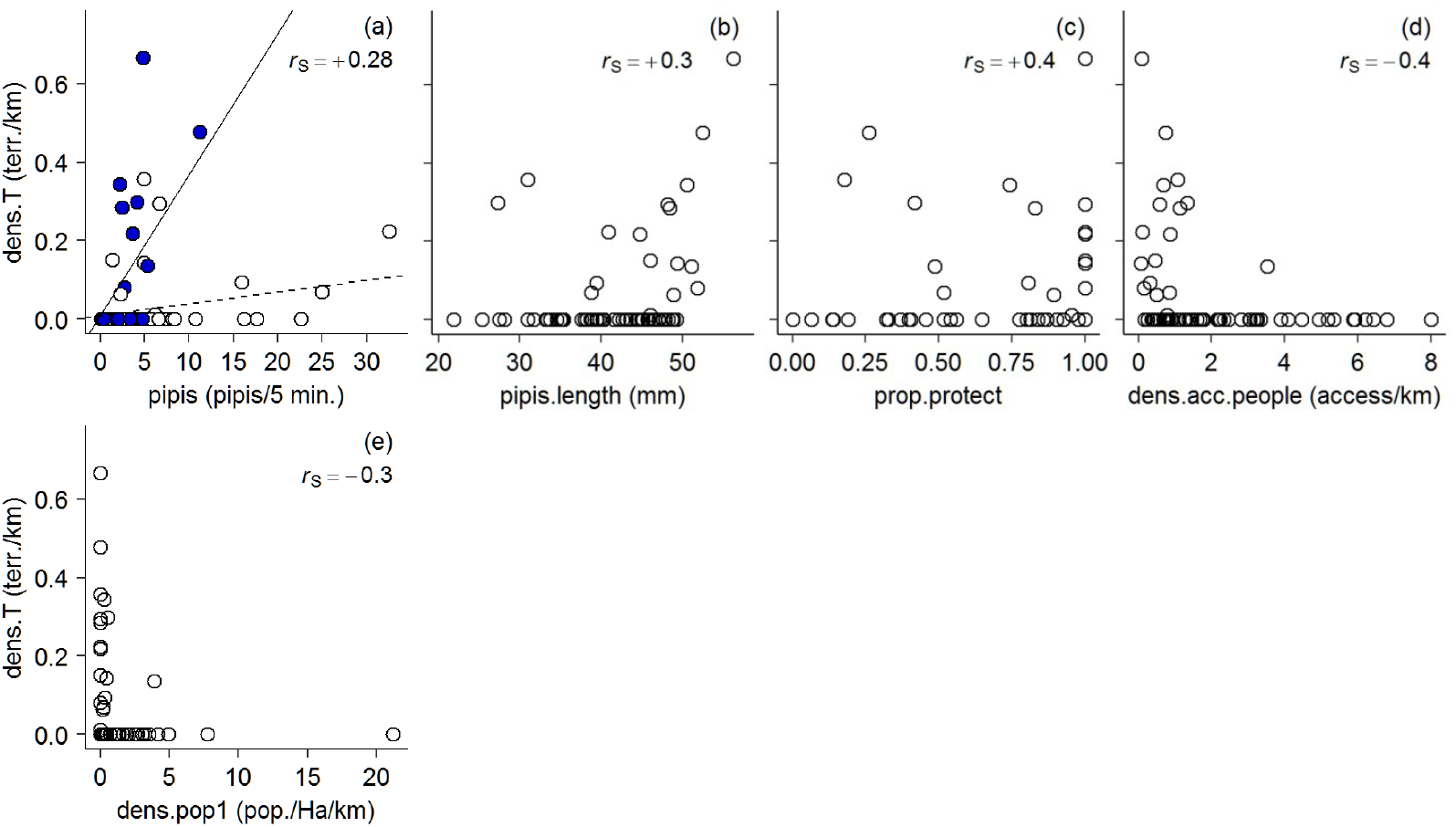
Oystercatcher territory density correlations.

Twelve beaches were sampled where commercial pipi fishing was observed or otherwise known to occur and 60 beaches without. Mean pipi count, mean pipi length, mean oystercatcher count density and mean oystercatcher territory density were greater for fished beaches than for non-fished beaches (Figure 6).

**Fig. 6.**
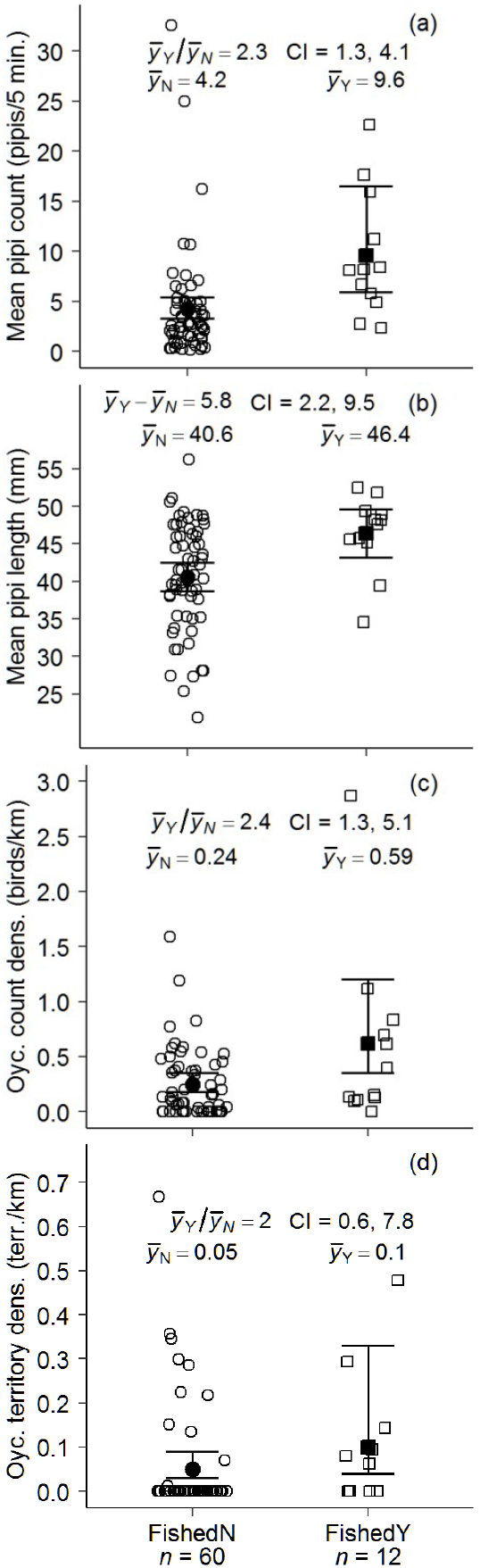
Pipi and oystercatcher contrasts for beaches with (squares) and without (circles) commercial pipi fishing. Individual results are plotted with a small amount of random horizontal ‘jitter’. Black points are means and error bars are 95% confidence intervals from generalised linear models (a, c, d) and an unequal variance *t*-test with Welch’s approximate degrees of freedom (b). Contrasts (mean ratios/difference) are from the same models and *t*-test.

### Multicollinearity

Hierarchical clustering of continuous input variables indicated three groups of collinear variables with Spearman *r*^2^ ≥ 0.49 (Fig. 7). The first cluster contained four coastal development and recreation variables. Mean people and dog count densities were dropped because those counts are imprecise, varying with the time of day, during the week, with weather conditions, *etc*. Oystercatcher count densities were equally correlated with the proportion of urban beach and human population density (Table 3) and those two variables were examined separately in further analyses. The second cluster contained spatial co-ordinates. Oystercatcher count densities were weakly correlated with UTM x and UTM y (Spearman *r* ≤ 0.29). UTM y was selected to account for possible latitudinal trends. The third cluster contained three vehicle-based recreation variables. Oystercatcher count densities were more strongly correlated with the proportion of 4WDs permitted beach (Spearman *r* ≥ 0.22) than with either of the other two variables, which were dropped.

**Fig. 7.**
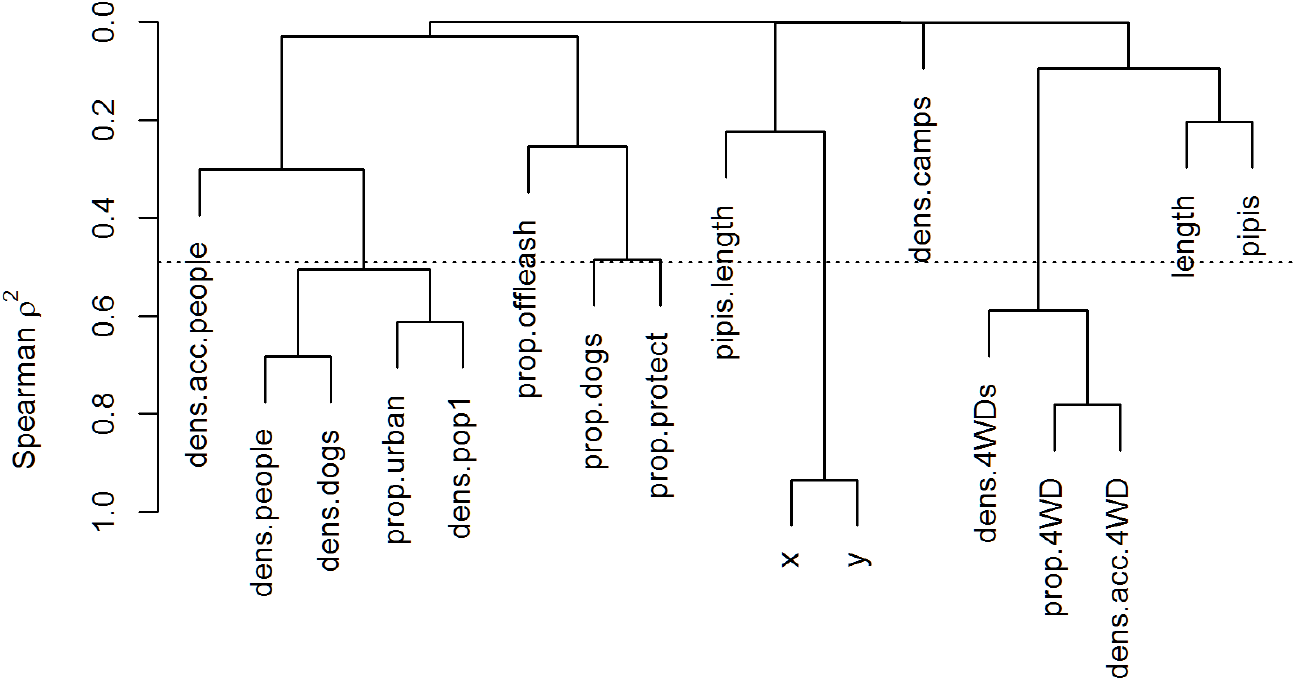
Hierarchical clustering of continuous input variables using squared Spearman rank-based correlations (Table 3). Variables are described in Table 2. The horizontal line cuts the dendrogram at Spearman *r*^2^ = 0.49 and indicates three clusters of collinear variables below the line: 1) *dens*.*people, dens*.*dogs, prop*.*urban, dens*.*pop1*; 2) *x, y*, and; 3) *dens*.*4WDs, prop*.*4WD, dens*.*acc*.*4WD*.

Next, large variance inflation factors resulted in UTM y (Spearman *r* = 0.03, VIF = 4.7) being dropped from mean oystercatcher count models and UTM y (Spearman *r* = 0.25, VIF = 6.2), camps and caravan parks count density (Spearman *r* = −0.01, VIF = 5.1), proportion of protected beach (Spearman *r* = 0.47, VIF = 3.1) and proportion of dogs permitted beach (Spearman *r* = −0.39, VIF = 2.5) being dropped from territory count models, in that order. The last two variables were dropped because they contributed to multicollinearity for the region predictor that was thought to be a necessary covariate for mean pipi count (Figs. 4a, 5a). After acting to reduce collinearity, the 19 input variables were reduced to 13 predictors for mean oystercatcher count and 10 for territory count (Table 2).

### Oystercatcher count models

Major features of the mean oystercatcher count ΔAICc < 6 model set were positive mean pipi count (except in model seven) and Far North Coast (*region2*OTH = 0) effects and a negative proportion of urban beach effect (or human population density in model six) (Table 4). There was no support for the pipi abundance null model (ΔAICc = 23.6; Fig. 8a). Including the proportion of urban beach improved the fit to lower densities (Figs. 8b,c).

**Table 4.**
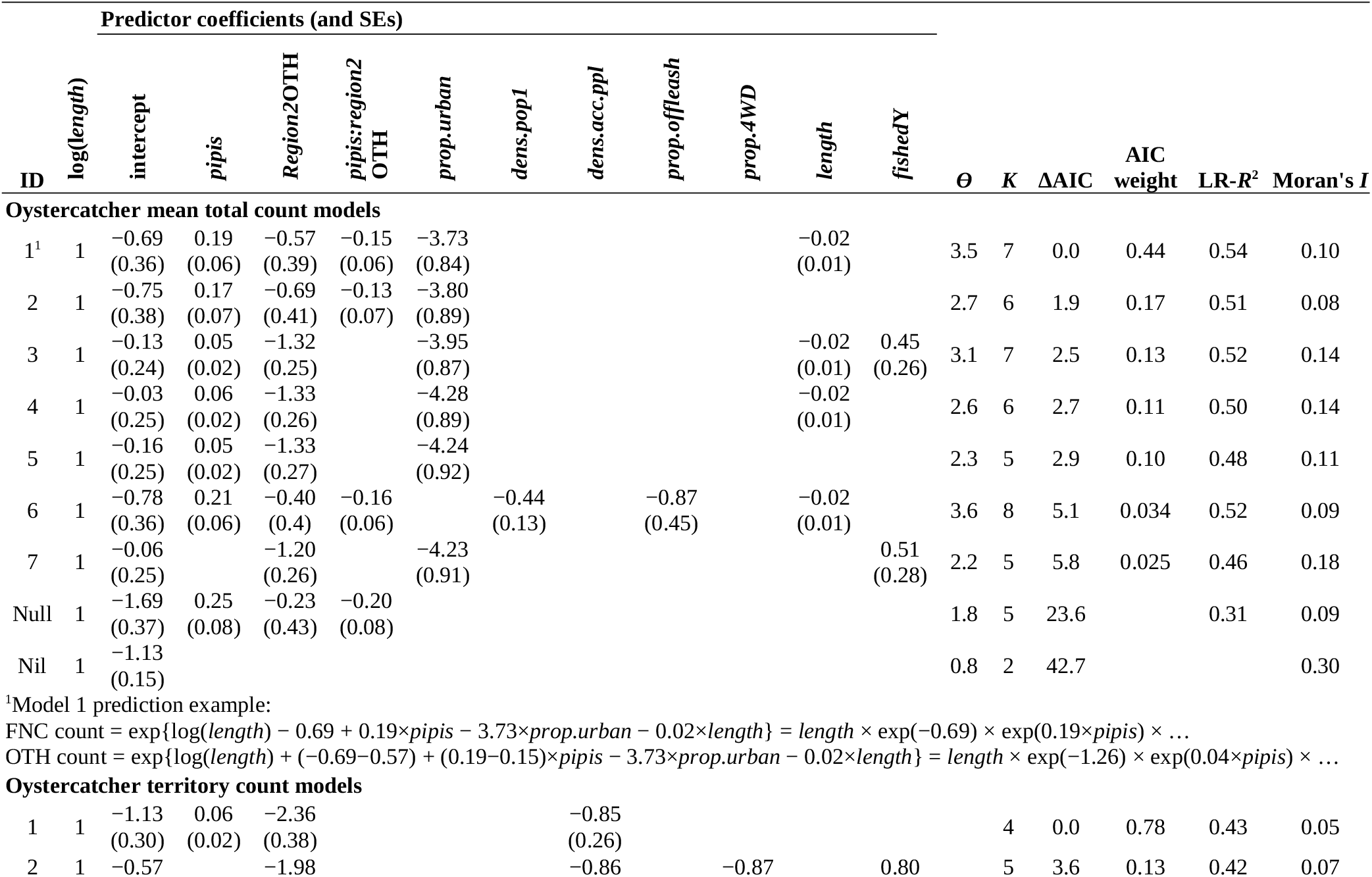
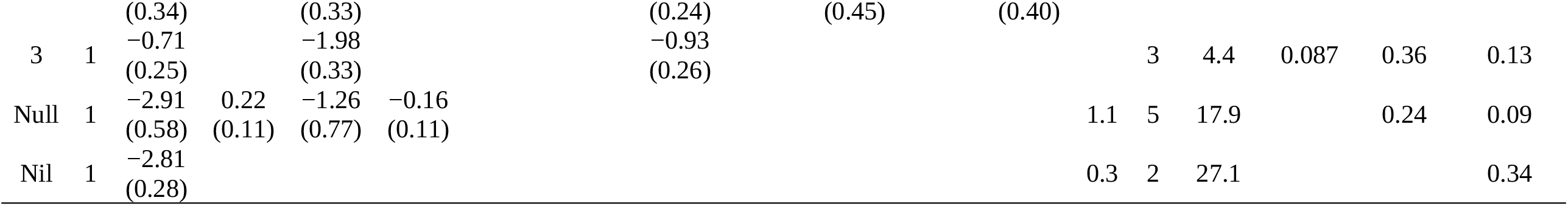
Summary of oystercatcher count generalised linear models sorted by small sample Akaike’s Information Criterion (AICc). ΔAICc = 0 indicates the ‘best approximating model’ of those in the model set; ΔAICc < 2 indicates strong support; ΔAICc = 2–7 indicates some support and ΔAICc > 14 indicates no support (Burnham *et al*. 2011). LR-*R*^2^ is likelihood-ratio pseudo-*R*^2^ (with reference to Nil models) and Moran’s *I* is an index of residual spatial autocorrelation. *ϑ* is the negative binomial overdispersion parameter (variance = mean + mean^2^/*ϑ*; blank cells indicate a Poisson error distribution), *K* equals the number of parameters in the model (number of predictors plus intercept plus ϑ). Predictors are described in Table 2 (blank cells indicate that a model did not include that predictor). The *region2* predictor is a binary factor with reference Far North Coast (OTH = other) and *region2*OTH and *pipis*:*region2*OTH are intercept and slope contrasts respectively. The *fished* predictor is a binary factor with reference N (Y = yes). The parameter log(*length*) is an offset.

**Fig. 8.**
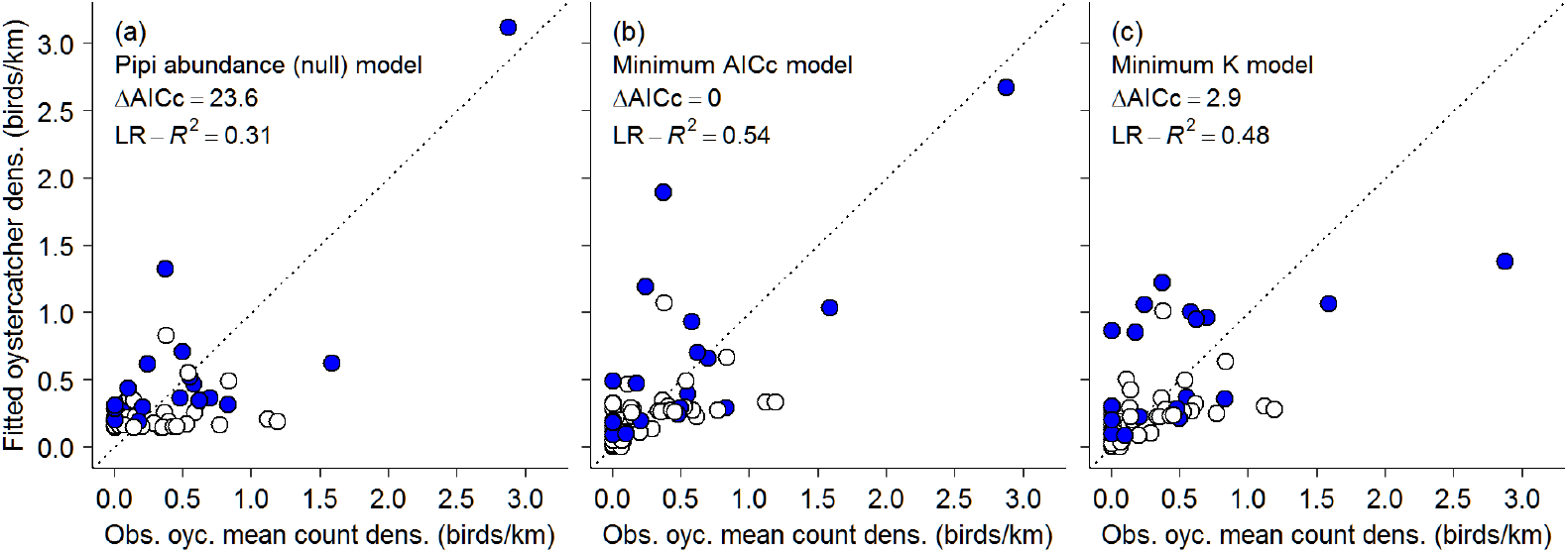
Fitted versus observed mean oystercatcher count density for the null (a) minimum AICc (b) and minimum *K* (simplest, model five) (c) models in Table 4. Far North Coast beaches (Coffs Harbour to the NSW-Queensland border) are plotted as blue-filled circles and other regions are plotted as white-filled circles.

Major features of the oystercatcher territory count ΔAICc < 6 model set were a positive Far North Coast effect and a negative pedestrian access count density effect (Table 4). There was no support for the pipi abundance null model (ΔAICc = 17.9; Fig 9a). Including pedestrian access density improved the fit but there were some false positives (observed = 0 and fitted > 0) and underestimates from the selected models (Figs 9b,c).

**Fig. 9.**
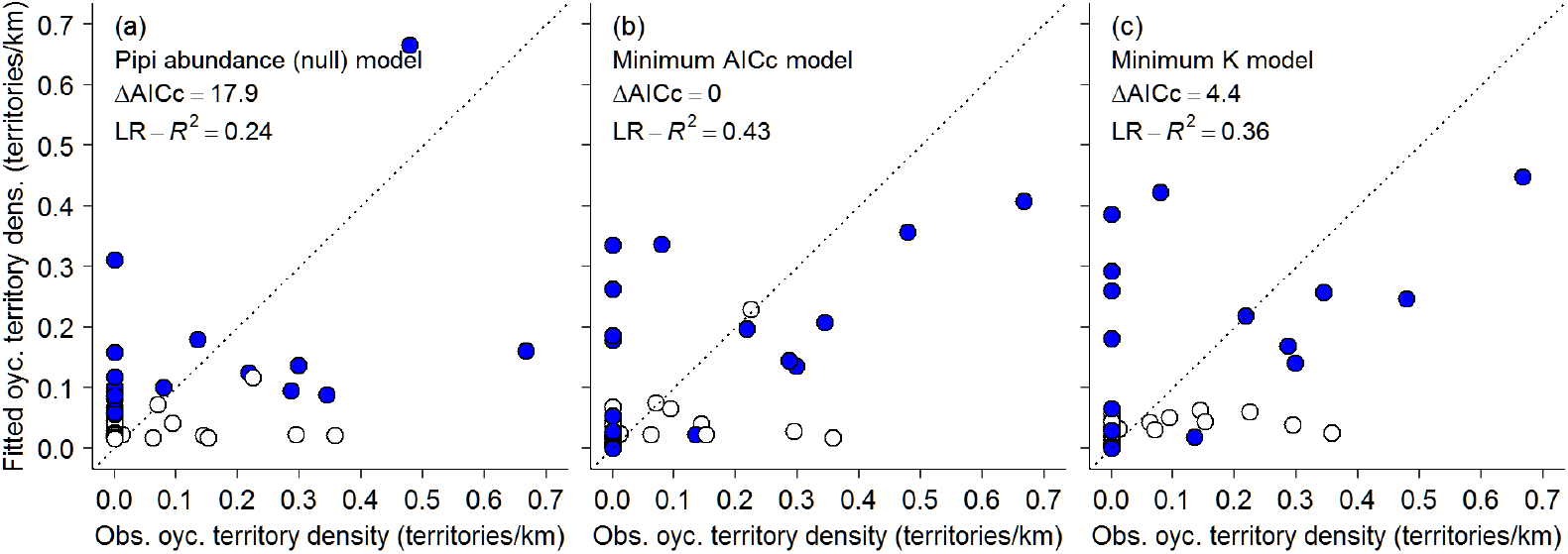
Fitted versus observed oystercatcher territory density for the null (a) minimum AICc (b) and minimum *K* (simplest, model four) (c) models in Table 4.

## DISCUSSION

The statistical analysis in this study was exploratory and descriptive because there was not the detailed ecological knowledge for Australian Pied Oystercatchers on ocean beaches for explanatory models (*e*.*g*. Goss-Custard 1996). Moreover, science can become ‘stuck’ in paradigms and exploratory, hypothesis-generating studies are necessary for science to evolve (Kuhn 1962, Smith 2000). With the usual caveat that ‘association is not causation’, the oystercatcher-habitat models presented suggest some important variables for conservation management and further research.

Oystercatcher counts in this study agreed with previous counts and estimates. The NSW Wader Study Group counted 232 oystercatchers on NSW ocean beaches in Oct 1998 (Owner & Rohweder 2003). There were 222 oystercatchers counted on NSW beaches in this study. The NSW Government has estimated that there are < 200 Australian Pied Oystercatcher breeding pairs in NSW (NSW DPIE 2019b), with 88 pairs at 15 Key Management Sites including 24 pairs at four beach sites: South Ballina, Broadwater (which was defined for management purposes as Airforce Beach N to Boundary Creek, at the S end of South Ballina Beach, Table 1), Bombing Range and a number of beaches within Yuraygir National Park (NSW DPIE 2019a). It is estimated that approximately one quarter (24/88 = 0.27) of oystercatcher pairs are beach-residents from which it can be estimated that there are < 54 (0.27 × 200) beach-resident pairs in NSW. There were 40 oystercatcher territories counted on NSW beaches in this study. A previous study counted 26 oystercatcher territories on beaches between the Richmond River and Bonville Creek (8 km S of Coffs Harbour) in 2003 and 27 in 2005 (Harrison 2009). There were 27 territories counted on Far North Coast beaches (N of Coffs Harbour) in this study. The agreement between these counts and estimates indicates that the 68 NSW beaches sampled in this study represented almost the entire state population of beach-resident oystercatchers.

Oystercatcher-habitat correlations were weaker than like correlations from the preceding study that sampled 11 Far North Coast beaches, N of the Clarence River (Owner & Rohweder 2003). This disagreement was largely resolved by recomputing correlations for only Far North Coast beaches (Table 5) however the strong oystercatcher-pipi correlation was then defined by only two beaches: South Ballina and Bombing Range (Fig 4a). There were several long and largely undeveloped Mid-north Coast beaches in this study with high mean pipi counts but only moderate oystercatcher count densities. Established pipi fisheries in both of these regions indicate that they both have a long history of high pipi abundance and, several years after the 2003–2009 ‘pipi crash’ (Harrison 2009, Totterman 2018), results from this ‘snap shot’ study are assumed to be representative of typical pipi abundances.

**Table 5.** Mean oystercatcher count density Pearson correlations compared with those from the preceding study that sampled 11 Far North Coast beaches, N of the Clarence River (Owner 1997, for birds/km and Owner & Rohweder 2003, for birds/km^2^). Far North Coast (FNC) beaches are those between Coffs Harbour and the NSW-Queensland border (Table 1, Fig. 1).

|  | Owner | Owner & | This study | This study |
| --- | --- | --- | --- | --- |
|  | (1997) | Rohweder | – | – |
| Mean oystercatcher |  | (2003) | all beaches | NSW FNC |
| count density ~ | *r* | *r* | *r* | *r* |
| Pipi mean density/count | 0.78 | 0.83 | 0.26 | 0.76 |
| Mean pipi length | 0.70 | 0.66 | 0.22 | 0.44 |
| Beach length | 0.55 | 0.64 | 0.09 | 0.72 |
| 4WD mean count density | 0.39 | 0.24 | −0.02 | 0.04 |
| People mean count density | −0.17 | −0.03 | −0.15 | −0.23 |

Automated information-theoretic model selection methods were effective at selecting small sets of reasonable models in this study. However, the usual rule of ‘garbage in, garbage out’ applies. Multicollinearity was particularly troublesome for territory count models, where oystercatcher-habitat associations were weak. Of the 19 input variables in this study, only four were frequent in the ΔAICc < 6 model sets for oystercatcher counts: mean pipi count, proportion of urban beach, region and pedestrian access count density. The proportion of urban beach and pedestrian access count density were not measured in previous oystercatcher-habitat studies. Alternatively, mean people count density was positively correlated with the proportion of urban beach in this study and could serve as a proxy for that variable. Mean people count density was measured in one of the previous studies, however the correlation with mean oystercatcher count density was weak (Owner & Rohweder 2003, Table 4). In the other previous study, distance to towns (isolation) was suggested to be a useful predictor for oystercatcher presence (Harrison 2009).

Mean oystercatcher count models in the ΔAICc < 6 set fitted observed count densities quite well. These models indicated that oystercatchers aggregate at Far North Coast beaches with high pipi abundance and that they avoid more urbanised beaches. Four of the seven models in the mean oystercatcher count ΔAICc < 6 model set included the region predictor but not as a covariate for mean pipi count. Model seven in that set did not include mean pipi count. These models provide further evidence that high oystercatcher count densities on the Far North Coast were a regional effect and, unlike what was proposed in previous studies (Owner & Rohweder 2003, Harrison 2009), not simply a predator-prey numerical response. High oystercatcher territory counts contributed to this Far North Coast effect and so local population histories and recruitment are suggested to be important. Proportion of urban beach (or human population density) substantially improved model fit for low mean oystercatcher count density beaches. The proposed explanation is that coastal development and associated human recreation disturbance displace shorebirds from otherwise suitable habitat (reviewed by Hockin *et al*. 1992, also see Harrison 2009).

Oystercatcher territory count models in the ΔAICc < 6 set fitted observed count densities rather poorly. The proposed explanation is that territorial birds are less responsive to habitat than are non-territorial birds. For example, pipi abundance at South Ballina declined to near zero during the 2003–2009 pipi crash but there remained eight territories and five breeding pairs on the beach in 2009 (Totterman 2018, 2020). Nonetheless, models in the ΔAICc < 6 set did indicate that territory density generally decreases towards zero on beaches with higher pedestrian access count density. The proposed explanation is that higher access density can result in shorter separation distances between access points and shorebird habitat zones, higher intrusion rates into those zones and more frequent disturbance (Dowling & Weston 1999, Totterman 2019c). With the exception of Tallow Beach, where spot mapping suggested one territory at Tallow Creek but where breeding is not known to occur, zero oystercatcher territories were observed in this study when pedestrian access density exceeded 1.3 tracks/km (Fig. 5d).

Beach access density is not only an issue for more urban beaches. Coastal tourism can also reduce the breeding range, density and breeding success for shorebirds (reviewed by Pienkowski 1993). More than two decades ago, it was proposed that tourism on Fraser Island was negatively impacting beach-nesting birds (Fisher *et al*. 1998). Teewah Beach and 75 Mile Beach (Fraser Island) in this study were long, largely undeveloped (both are within Great Sandy National Park) and had high mean pipis counts but also extensive beach front camping zones with high count densities of pedestrian and 4WD tracks. Spot mapping indicated only one oystercatcher territory for these two beaches with a combined length of 132 km. Moreover, mean oystercatcher counts on 75 Mile Beach have declined from 15 birds between Hook Point and Indian Head in 1995 (Fisher *et al*. 1998) to three in this study (Table 1). Together with the negative proportion of urban beach effect for mean oystercatcher counts, it is recommended that the coastal development and human recreation disturbance threats for the Australian Pied Oystercatcher be upgraded.

No overall negative fishing effects for pipi stocks and oystercatcher abundance were found in this study. Instead, mean pipi biomass and mean oystercatcher count and territory count densities were higher on commercially fished beaches than on non-fished beaches. These results suggest that both fishers and oystercatchers target beaches with higher pipi biomass. The purposive sampling design in this study selected beaches where pipis and oystercatchers could be abundant, including all known fished beaches. The sample of fished beaches is assumed to be exhaustive and positive bias is assumed for pipi and oystercatcher means for the non-fished beach sample (which would act to reduce, rather than increase, observed differences). Together with uncertainty about the oystercatcher-pipi response, it is recommended that the pipi harvesting threat for the Australian Pied Oystercatcher be downgraded.

Coastal development and human recreation disturbance appeared to limit oystercatcher population sizes on E Australian beaches in this study. With commercial interests for developing coastal land and social interests for maintaining the *status quo* with regard to access and recreation (James 2000, Priest *et al*. 2002), degradation and loss of beach-nesting bird habitat is set to continue. Conservation efforts for Australian Pied Oystercatchers in NSW, which have been almost exclusively focussed on breeding success (Wellman *et al*. 2000, NSW Office of Environment & Heritage 2011), will fail in the long term unless important habitat for the species is recognised and protected from development and intense human recreation (*e*.*g*. Totterman 2020).

## ACKNOWLEDGEMENTS

This paper was submitted to Stilt, peer-reviewed and then ‘dropped’ silently, having not been published to-date.

## REFERENCES

Albert, C.H., N.G. Yoccoz, T.C. Edwards Jr, C.H. Graham, N.E. Zimmermann & W. Thuiller. 2010. Sampling in ecology and evolution – bridging the gap between theory and practice. Ecography 33: 1028–1037. https://doi.org/10.1111/j.1600-0587.2010.06421.x

Arnold, T.W. 2010. Uninformative parameters and model selection using Akaike’s Information Criterion. Journal of Wildlife Management 74: 1175–1178. DOI: 10.2193/2009-367

Australian Bureau of Statistics. 2011. Australian Population Grid 2011. Australian Government, ABS. Accessed 28 May 2017 at: https://www.abs.gov.au/ausstats/abs@.nsf/mf/1270.0.55.007

Barton, K. 2020. MuMIn: Multi-Model Inference. R package version 1.43.17. https://CRAN.R-project.org/package=MuMIn

Bivand, R.S, E. Pebesma &V. Gomez-Rubio. 2013. Applied spatial data analysis with R, Second edition. Springer, New York, U.S.A.

Bivand, R., T. Keitt & B. Rowlingson. 2021. rgdal: Bindings for the ‘Geospatial’ Data Abstraction Library. R package version 1.5-23. https://CRAN.R-project.org/package=rgdal

Bjørnstad, O.N & W. Falck. 2001. Nonparametric spatial covariance functions: Estimation and testing. Environmental and Ecological Statistics 8: 53–70.

Bjørnstad, O.N. 2020. ncf: Spatial Covariance Functions. R package version 1.2-9. https://CRAN.R-project.org/package=ncf

Burnham, K.P., D.R. Anderson & K.P. Huyvaert. 2011. AIC model selection and multimodel inference in behavioral ecology: some background, observations, and comparisons. Behavioral Ecology and Sociobiology 65: 23–35.

Cade, B.S. 2015. Model averaging and muddled multimodel inferences. Ecology 96: 2370–2382.

Dormann, C.F., J. Elith, S. Bacher, C. Buchmann, G. Carl, G. Carré, J.R. García Marquéz, B. Gruber, B. Lafourcade, P.J. Leitão, T. Münkemüller, C. McClean, P.E. Osborne, B. Reineking, B. Schröder, A.K. Skidmore, D. Zurell & S. Lautenbach. 2013. Collinearity: a review of methods to deal with it and a simulation study evaluating their performance. Ecography 36: 27–46. doi: 10.1111/j.1600-0587.2012.07348.x

Dowling, B. & M.A. Weston. 1999. Managing a breeding population of the Hooded Plover Thinornis rubricollis in a high-use recreational environment. Bird Conservation International 9: 255–270.

Dunn, P.K. & G.K. Smyth. 1996. Randomized quantile residuals. Journal of Computational and Graphical Statistics 5: 236–244.

Ferguson, G., D. Johnson & H. Gorfine. 2018. Pipi Donax deltoides. In: Status of Australian Fish Stocks Reports 2018 (C. Stewardson, J. Andrews, C. Ashby, M. Haddon, K. Hartmann, P. Hone, P. Horvat, S. Mayfield, A. Roelofs, K. Sainsbury, T. Saunders, J. Stewart, S. Nicol & B. Wise, Eds.). Fisheries Research and Development Corporation, Canberra, Australia. http://www.fish.gov.au/report/262-Pipi-2018

Fisher, F., M. Hockings & R. Hobson. 1998. Recreational impacts on waders on Fraser Island. Sunbird 28: 1–11.

Fox, J. & S. Weisberg. 2019. An {R} Companion to Applied Regression, Third Edition. Sage, Thousand Oaks, California, USA.

Fraser, N. & Lindsey, A. 2018. Some observations of Australian Pied Oystercatcher on Worimi Conservation Lands. Whistler 12: 35–42.

Gelman, A. & J. Hill. 2007. Data Analysis Using Regression and Multilevel/Hierarchical Models. Cambridge UniversityPress, Cambridge, UK.

Goss-Custard, J.D., Ed. 1996. The oystercatcher: from individuals to populations. Oxford University Press, Oxford, UK.

Graham, M.H. 2003. Confronting multicollinearity in ecological multiple regression. Ecology 84: 2809–2815.

Gray, C.A. 2016a. Effects of fishing and fishing closures on beach clams: experimental evaluation across commercially fished and non-fished beaches before and during harvesting. PLoS ONE 11: e0146122. https://doi.org/10.1371/journal.pone.0146122.

Gray, C.A. 2016b. Evaluation of fishery-dependent sampling strategies for monitoring a small-scale beach clam fishery. Fisheries Research 177: 24–30.

Harrell, F.E. (with contributions from C. Dupont and many others). 2021. Hmisc: Harrell Miscellaneous. R package version 4.5-0. https://CRAN.R-project.org/package=Hmisc

Harrison, A.E. 2009. The ecology of two vulnerable shorebirds (Haematopus fuliginosus and H. longirostris) in sub-tropical northern NSW, Australia: implications for conservation and management. PhD thesis, University of New England, Armidale, Australia.

Hijmans, R.J. 2020. raster: Geographic Data Analysis and Modeling. R package version 3.4-5. https://CRAN.R-project.org/package=raster

Hockin, D., M. Ounsted, M. Gorman, D. Hill, V. Keller & M.A. Barker. 1992. Examination of the effects of disturbance on birds with reference to its importance in ecological assessments. Journal of Environmental Management 36: 253–286.

James, R. J. 2000. From beaches to beach environments: linking the ecology, human-use and management of beaches in Australia. Ocean and Coastal Management 43: 495–514.

Kuhn, T.S. 1962. The Structure of Scientific Revolutions. First Edition. University of Chicago Press, Chigao, USA.

Marchant S. & P.J. Higgins, Eds. 1993. Haematopus longirostris Pied Oystercatcher. PAges 716–726 in: Handbook of Australian, New Zealand and Antarctic birds. Vol. 2, raptors to lapwings. Oxford University Press, Melbourne, Australia.

Murray-Jones, S. 1999. Conservation and management in variable environments: the surf clam, Donax deltoides. PhD thesis, University of Wollongong, Australia.

Murtaugh, P.A. 2007. Simplicity and complexity in ecological data analysis. Ecology 88: 56–62.

Nagelkerke, N.J.D. 1991. A note on a general definition of the coefficient of determination. Biometrika 78: 691–692.

NSW Department of Planning, Industry and Environment. 2019a. Saving our species. Pied Oystercatcher (Haematopus longirostris) Key Management Sites. NSW Government, DPIE. Accessed 26 Aug 2019 at: https://www.environment.nsw.gov.au/savingourspeciesapp/project.aspx?ProfileID=10386

NSW Department of Planning, Industry and Environment. 2019b. Threatened species. Pied Oystercatcher - profile. NSW Government, DPIE. Accessed 26 Aug 2019 at: https://www.environment.nsw.gov.au/threatenedSpeciesApp/profile.aspx?id=10386

NSW Government. 2016. Biodiversity Conservation Act 2016, No. 63. Accessed 18 Feb 2019 at: https://www.legislation.nsw.gov.au/~/view/act/2016/63/

NSW Office of Environment & Heritage. 2011. NSW Threat abatement plan for predation by the red fox (Vulpes vulpes). NSW Office of Environment and Heritage, Sydney. Accessed 13 May 2020 at: https://www.environment.nsw.gov.au/resources/pestsweeds/110791FoxTAP2010.pdf

NSW Scientific Committee. 2010. Pied Oystercatcher Haematopus longirostris Vieillot 1817 -endangered species listing, final determination. Accessed 11 Nov 2018 at: https://www.environment.nsw.gov.au/determinations/piedoystercatcherFD.htm

Owner, D. 1997. The ecology and management of the Pied Oystercatcher (Haematopus longirostris) in Northern NSW. BSc thesis, Southern Cross University, Lismore, Australia.

Owner, D. & D.A. Rohweder. 2003. Distribution and habitat of Pied Oystercatchers (Haematopus longirostris) inhabiting beaches in northern New South Wales. Emu 103: 163–170.

Pienkowski, M.W. 1993. The impact of tourism on coastal breeding waders in western and southern European overview. Wader Study Group Bulletin 68: 92–96.

Priest, B., P. Straw & M.A. Weston. 2002. Shorebird conservation in Australia. Wingspan 12 (Supplement): 1–15.

R Core Team. 2021. R: A Language and Environment for Statistical Computing. Version 4.04. Vienna: R Foundation for Statistical Computing. http://www.R-project.org

Richards, S.A. 2008. Dealing with overdispersed count data in applied ecology. Journal of Applied Ecology 45: 218–227. https://doi.org/10.1111/j.1365-2664.2007.01377.x

Short, A.D. 2007. Beaches of the New South Wales coast. Second Edition. Sydney University Press, Sydney, Australia.

Smith, J. 2000. Nice work — but is it science? Nature 408: 293.

Totterman, S.L. 2018. Response of Australian Pied Oystercatchers Haematopus longirostris to increasing abundance of the beach bivalve prey Donax deltoides. Unpubl. preprint. bioRxiv. https://doi.org/10.1101/444786

Totterman, S.L. 2019a. Seasonal zonation patterns of the sandy beach bivalve Donax deltoides (Bivalvia: Donacidae) in subtropical eastern Australia. Unpubl. prepint. bioRxiv. https://doi.org/10.1101/610576

Totterman, S. 2019b. Distances walked by beach users and protecting shorebird habitat from human recreation disturbance. Unpubl. preprint. bioRxiv. https://doi.org/10.1101/783696

Totterman, S.L. 2019c. A “feet digging” swash zone sampling method for the sandy beach bivalve Donax deltoides (Bivalvia: Donacidae). Unpubl. preprint. bioRxiv. https://doi.org/10.1101/686196

Totterman, S. 2020. Taking fright: the decline of Australian Pied Oystercatchers Haematopus longirostris at South Ballina Beach, New South Wales. Unpubl. prepint. bioRxiv. https://doi.org/10.1101/2020.06.09.141796

Taylor, I.R. & S.G. Taylor. 2005. Foraging behaviour of Pied Oystercatchers in the presence of kleptoparasitic Pacific Gulls. Waterbirds 28: 156–161.

Taylor, I.R., O.M.G. Newman, P. Park, B. Hansen, C.D.T. Minton & R. Jessop. 2014. Conservation assessment of the Australian Pied Oystercatcher Haematopus longirostris. International Wader Studies 20: 116–128.

Venables, W.N. & B.D. Ripley. 2002. Modern Applied Statistics with S. Fourth Edition. Springer, New York, USA.

Ver Hoef, J.M. & Boveng, P.L. 2007. Quasi-poisson vs. negative binomial regression: how should we model overdispersed count data? Ecology 88: 2766–2772. https://doi.org/10.1890/07-0043.1

Warton, D.I. 2005. Many zeros does not mean zero inflation: comparing the goodness-of-fit of parametric models to multivariate abundance data. Environmetrics 16: 275–289.

Wellman L., B. Moffatt, B. Totterman & N. Hing. 2000. A co-operative approach to protecting threatened species - Pied Oystercatchers, South Ballina, Northern NSW. Pages 99– 104 in: Proceedings of the NSW pest animal control conference, 25–27 Oct 2000 (S. Bulogh, Ed.). NSW Agriculture, Orange.

Weston, M.A., E.M. McLeod, D.T. Blumstein & P.-J. Guay. 2012. A review of flight initiation distances and their application to managing disturbance to Australian birds. Emu 112: 269–286.

Zuur, A.F., E.N. Ieno & C.S. Elphick. 2010. A protocol for data exploration to avoid common statistical problems. Methods in Ecology and Evolution 1: 3–14. https://doi.org/10.1111/j.2041-210X.2009.00001.x

